# Wnt5A Signaling Blocks Progression of Experimental Visceral Leishmaniasis

**DOI:** 10.1101/2021.11.12.468361

**Authors:** Shreyasi Maity, Arijit Chakraborty, Sushil Mahata, Syamal Roy, Anjan Kumar Das, Malini Sen

## Abstract

Visceral Leishmaniasis, caused by *L. donovani* infection is fatal if left untreated. The intrinsic complexity of visceral leishmaniasis complicated further by the increasing emergence of drug resistant *L. donovani* strains warrants fresh investigations into host defense schemes that counter infections. Accordingly, using a mouse model of experimental visceral leishmaniasis we explored the utility of host Wnt5A in restraining *L. donovani* infection, using both antimony sensitive and antimony resistant *L. donovani* strains. We found that Wnt5A heterozygous (Wnt5A +/-) mice are more susceptible to *L. donovani* infection than their wild type (Wnt5A +/+) counterparts as depicted by the respective Leishman Donovan Units (LDU) enumerated from the liver and spleen harvested from infected mice. Higher LDU in Wnt5A +/-mice correlated with increased level of plasma gammaglobulin, liver granuloma and disorganization of splenic germinal centers. Progression of infection in mice by both antimony sensitive and antimony resistant strains of *L. donovani* could be prevented by activation of Wnt5A signaling as evident from the lowered LDU and gammaglobulin level, and intactness of splenic germinal centers through intravenous administration of rWnt5A prior to *L. donovani* infection. Wnt5A mediated blockade of *L. donovani* infection correlated with the preservation of splenic macrophages and activated T cells, and a TH1 like cytokine thrust. Taken together our results indicate that depletion of Wnt5A promotes susceptibility to visceral leishmaniasis and revamping Wnt5A signaling in the host is able to curb *L. donovani* infection irrespective of antimony sensitivity or resistance and mitigate the progression of visceral leishmaniasis.

## Introduction

Visceral Leishmaniasis (VL) or Kala-azar is a vector borne disease transmitted by the sand fly and caused by infection with the parasite *Leishmania donovani* (*L. donovani*). *L. donovani* promastigotes delivered to the host during a blood meal by the sand fly differentiate within host macrophages to amastigotes, which replicate in macrophages and build their niche therein at the cost of the host cell machinery. Globally, VL is among the top ten neglected tropical diseases in the World Health Organization list. There are outbreaks of VL in certain pockets in developing countries and the disease if left untreated can turn out fatal [(1–3), https://www.who.int/news-room/fact-sheets/detail/leishmaniasis]. Although different modes of therapy for VL are available, the increasing emergence of drug resistant strains makes treatment complicated (3–5). In this scenario it is important to understand the intricacies of the host – parasite interactions and how host immunity can be boosted to counter the parasite infection. Keeping in mind the documented role of Wnt5A signaling in the regulation of bacterial infections and immune homeostasis (6–10), we became interested in evaluating the influence of Wnt5A on *L. donovani* infection and progression of VL.

Wnt5A, a secreted glycoprotein belongs to a 19 - member family of Wnt ligands that interact with Frizzled and/or ROR cell surface receptors during signal transduction in cells. On account of considerable homology among the Wnts and the Wnt receptors, overlap exists in the pairing of Wnts with their cognate receptors leading to cross talk among the intracellular intermediates at the different levels of Wnt signaling. Classically, the transcriptional coactivator β-catenin is essential for the canonical mode of Wnt signaling, but in the non-canonical mode of Wnt signaling represented by Wnt5A, β-catenin is dispensable (11–16).

Wnt5A signaling sustains cell differentiation and polarity. This property of Wnt5A signaling is utilized in the maintenance of immune homeostasis, wherein it regulates cytoskeletal actin dynamics during macrophage mediated phagocytosis of microbes and their autophagic clearance (8,10,17,18). In the context of *L. donovani* infection, we previously demonstrated that a Wnt5A-Rac1-Actin axis prevents the growth of *L. donovani* in macrophages by abolishing the existence of parasitophorous vacuoles through enhancement of prasitophorous vacuole – lysosome fusions (19). These studies prompted us to investigate the potential of Wnt5A in restraining experimental visceral leishmaniasis caused by *L. donovani* infection in mice.

In the current study we have demonstrated that genetic depletion of Wnt5A in Wnt5A heterozygous mice leads to increased susceptibility to the development and progression of disease upon *L. donovani* infection. Administration of recombinant Wnt5A (rWnt5A) prior to *L. donovani* infection, on the other hand, makes mice resistant to the development of experimental visceral leishmaniasis. Blockade in disease pathogenesis by Wnt5A signaling may be attributed to altered T cell and macrophage activation and regulation of cytokine expression.

## Materials and Methods

### Reagents

Medium used for cell and parasite culture was RPMI 1640 (Cat No. -31800-022) and Medium 199 (Cat No.-11150-059) were purchased from Gibco, USA. Antibodies, APC Rat anti-mouse CD138 (Cat No. - 558626), V500 Syrian Hamster anti-mouse CD3 (Cat No. - 560771), PerCP-Cy5.5 Rat Anti-mouse CD4 (Cat No. - 550954), BV605 Rat Anti-mouse CD8 (Cat No. - 563152) and PE-CF594 Rat Anti-mouse IFN-γ were purchased from BD Biosciences, PE-Cyanine 7 Rat anti-mouse Granzyme B (Cat. No.- 25-8898-82), anti-mouse CD169 (Cat no.-MA516508), anti-mouse CD209b (Cat no. – 14-2093-82), goat anti-rat IgG secondary alexa fluor 488 (Cat no.-A-11006), goat anti-hamster IgG alexa fluor 647 (Cat no. - A-21451) were purchased from eBioscience, anti-mouse F4/80 was purchased from Santa Cruz, Anti-Human/Mouse Wnt5A monoclonal antibody (Cat no.- MAB645) and anti-Rat IgG-HRP (Cat no.- HAF005) were purchased from R & D Systems, anti-human IL-6 antibody (Cat no.- BB-AB0150) was purchased from Biobharati Life Science. Anti-rabbit IgG-HRP (Cat no.- A0545) and Anti-mouse IgG-HRP (Cat no.- A9044) were purchased from Sigma. Anti-mouse IL-10 ELISA set (Cat no. – 555252), anti-mouse IFN-γ ELISA set (Cat no. – 555138) and anti-human IL-10 ELISA set (Cat no. – 555157) were purchased from BD Biosciences. Fetal Bovine Serum (Cat no. – 10082147), PenStrep (Cat no.- 15140122), L-Glutamine (Cat no. – 25030081), DAPI (Cat no.-D1306), alexa fluor 555 phalloidin (Cat no. – A34055) and cell tracker green (Cat no.- C2925) were purchased from Invitrogen. Histopaque solution (Cat no.- 10771-100ml) and poly-l-lysine (Cat no.- P-8920) were purchased from Sigma. TMB solution (Cat no.- CL07-1000MLCN) was purchased from Merck. Bradford reagent (Cat no. -5000006) was purchased from Biorad, USA. Recombinant Wnt5A (Cat no.- 10UG 645-WN-010 R & D, GF146 Millipore) and recombinant Wnt3A (Cat no.- 10UG 5036-WN-010 R & D, GF154 Millipore) were purchased from R & D Systems and Millipore. BSA (SRL-83803), DCF-DA (Calbiochem CAS 4091-99-0)

### Animal Maintenance

BALB/c mice were maintained in institute animal facility. 129S7-*Wnt5Atm1Amc*/J mice were purchased from Jackson Laboratory, USA and breeding was done in in-house animal facility. Genotyping was done to differentiate Wnt5A wild type from Wnt5A heterozygous mice by following Jackson laboratory protocol [(8),https://www.jax.org/Protocol?stockNumber=004758&protocolID=23556]. Animals were kept in IVC <u>(I</u>ndividually <u>V</u>entilated <u>C</u>aging) system with ad libitum water and food under optimum physiological temperature and balanced light and dark cycle. All experimental groups comprised mice of 8-12 weeks old.

### Parasite and Cell Maintenance

*L. donovani* WHO reference strain AG83 [MHOM/IN/1983/AG83] and Stibogluconate resistant strain BHU575 [MHOM/IN/2009/BHU575/0] were regenerated from BALB/c mice infected separately with the two different strains. Promastigote form of the parasite was maintained in a B.O.D. shaker incubator at 22°C in Medium 199 supplemented with 20% FBS, 2 mM L-glutamine, 100 U/ml penicillin, and 100mg/ml streptomycin. Freshly transformed stationary-phase promastigotes were harvested and washed in 0.02 M phosphate buffered saline, pH 7.2 for injection into mice. The BHU575 strain was a kind gift from Dr. Shyam Sundar, BHU, Varanasi, India.

Peritoneal macrophages were isolated from BALB/c mice. The peritoneum was washed with ice-cold PBS for harvesting peritoneal lavage. Then the lavage was centrifuged at 2000 rpm and the peritoneal cells were resuspended in RPMI 1640 medium supplemented with 10% FBS, 2 mM L-glutamine, 100 U/ml penicillin, and 100 mg/ml streptomycin.

### Quantification of parasite load in liver and spleen of animals

Each experimental mouse was infected with 10^8^ promastigotes through the tail vein and sacrificed either 45 days or 110 days post infection for generating imprints of the infected liver and spleen. For protein pretreatment experiments, each mouse was intravenously injected with 100ng rWnt5A on consecutive days before the infection and subsequent imprint collection. Tissue imprints were fixed in chilled absolute methanol and stained with Giemsa. Giemsa-stained micrographs were captured in the bright field of Leica microscope under 100X magnification with a Leica DFC450c camera. LDU (Leishman Donovan Unit) calculation was done based on at least 30 microscopic fields by counting the number of amastigotes per 1000 host nuclei and multiplying by respective organ weight in milligram (20).

### Tissue histology and histological scoring

Tissue histology was performed following published protocols (21). Mouse spleen and liver was dissected out and fixed in 10% buffered formalin overnight. Following dehydration in graded alcohol, each paraffin embedded tissue was sectioned using a microtome at a thickness of 3 µM, subsequent to which the sections were mounted in Mayer’s albumin-coated glass slides and stained with hematoxylin–eosin for microscopic analysis.

In H & E-stained spleen tissue sections histological scoring was done following previously published protocol with minor modifications (22). The degree of white pulp structural organization was evaluated based on 4 different criteria: well organized (A: distinct germinal centre and marginal zone), slightly disorganized (B: slight loss in distinctness of germinal center and marginal zone), moderately disorganized (C: poorly individualized or indistinct germinal center and marginal zone) and intensely disorganized (D: distinctness of white pulp from the red pulp area barely visible). In each case, the percent organization of the spleen in terms A, B, C & D was assessed by dividing the number of white pulps under each category by the total number of countable white pulp and multiplying the ratio by 100. For liver histological scoring, the total number of granulomas were counted from at least 20 microscopic fields for each experiment.

### Flow cytometry

Single cell suspensions generated from mouse spleen (teasing harvested spleen on mesh and collection of cells through strainer), and PBMC harvested from mouse cardiac blood (https://www.infinity.inserm.fr/wp-content/uploads/2018/01/PBMC-isolation-and-cryopreservation.pdf) were subjected to analysis by flow cytometry. Depending upon the purpose of the experiment, cells were surface stained separately for 1 hour at 4° C with different antibodies: anti-CD138-APC (for plasma cell), anti-F4/80-FITC (for macrophage), anti-CD169 and anti-rat Alexa fluor 488 (for marginal metallophillic macrophage), anti-CD3-HorizonV500 (for CD3), anti-CD4-PerCPCy5.5 (for CD4) and anti-CD8-BV605 (for CD8). Subsequently, cells were washed and processed for analysis. For both intracellular and surface staining, splenocytes were plated and incubated overnight under normal tissue culture conditions with addition of Brefeldin A (3µg/mL) before the last 4 hours of incubation. Subsequently, cells were fixed with 1% paraformaldehyde and permeabilized using permeabilization buffer (0.05% Tween 20 + 1% BSA in 1X PBS) before finally staining with anti-CD3-HorizonV500, anti-CD4-PerCPCy5.5, anti-CD8-BV605, anti-IFNγ-PE-CF594 and anti-Granzyme-B-PE-Cy7. Data acquisition was done in BD.LSR Fortessa Cell analyser. Data was analyzed using FCS express 5 software. Gating strategies are explained in the figures.

### Confocal Microscopy

Tissue sections, processed as described before were mounted on poly-L-lysine-coated glass slides for immunofluorescence microscopy following standard protocol (23,24). The prepared slides were immersed in sodium citrate buffer [10 mM sodium citrate, 0.05% Tween 20 (pH 6.0)] and placed in a pressure cooker for heat-induced antigen retrieval. The slides were then incubated with primary antibody (anti-mouse CD209b antibody) overnight at 4°C after blocking with 10% normal serum and 1% BSA in TBS (Tris Buffered Saline). A fluorophore-conjugated secondary antibody (anti-hamster 2° antibody alexa fluor 647 conjugated) was used to detect the signal under 40X magnification of TCS-SP8 confocal microscope.

Peritoneal exudate was collected from BALB/c mice and plated for 6hrs. The peritoneal macrophages were infected with cell tracker green stained *L. donovani* AG83 promastigotes for 4hrs at 5 MOI and washed 3 times with chilled 1X PBS, following which incubation was continued for 12 hrs in fresh culture medium. Subsequently the cells were fixed in 4% paraformaldehyde for 15 min, and Phalloidin and DAPI staining was done using Alexa Fluor 555 Phalloidin (1:2,000) and DAPI (1:4,000) PBST (0.1% Tween-20) with 2.5% BSA for 15 min. Following 3X PBST wash, the slides were mounted with 60% glycerol and visualized under 63X magnification (2.51 zoom) of TCS-SP8 confocal microscope.

### Transmission Electron Microscopy (TEM)

**Thin slices (~2 mm thick) of** *L. donovani* infected mouse spleen and liver were immersion fixed with 2% glutaraldehyde in 0.1 M cacodylate buffer and postfixed further with1% OsO_4_ in 0.1 M cacodylate buffer on ice for 1 hr. The fixed tissue was stained with 2% uranyl acetate for 1 hr on ice, after which it was dehydrated in graded series of ethanol (50-100%) while remaining on ice. The fixed dehydrated tissue was subjected to 1 wash with 100% ethanol and 2 washes with acetone (10 min each) and finally embedded with Durcupan. Subsequently, sections were cut at 60 nm on a Leica UCT ultramicrotome, and mounted on Formvar and 300 mesh copper grids. Subsequently, the sections were stained with 2% uranyl acetate for 5 minutes and Sato’s lead stain for 1 minute. Finally, grids were viewed using a JEOL JEM1400-plus TEM (JEOL, Peabody, MA) and photographed using a Gatan OneView digital camera with 4×4k resolution (Gatan, Pleasanton, CA).

### Mouse cytokine and anti-Leishmanial IgG estimation

IL-10 and IFN-γ of mouse plasma was measured with BD ELISA Set (catalog numbers 555138 and 555252 for IFN-γ and IL10 respectively) following manufacturers protocol (www.bdbiosciences.com). Briefly 96 well polystyrene ELISA plates were coated with capture antibody and kept at 4°C overnight. After washing with PBST, plates were blocked with 10% FBS in 1X PBS for 1 hour. Following the addition 100 μl of 1:10 diluted plasma samples, the plates were incubated with biotinylated detection antibody and streptavidin HRP. Subsequently TMB was added and Stop solution (2N H_2_SO_4_) was used after 30 minutes to stop the color reaction. Reading was taken in 450 nm in an ELISA plate reader.

Anti-Leishmanial IgG estimation was performed following previously published protocol (25). Briefly, stationary-phase promastigotes of *L. donovani* AG83 and *L. donovani* BHU575 were harvested for antigen preparation. After 4X wash with ice-cold PBS, the parasite pellet was suspended at a concentration of about 1.0 g of pellet in 50 ml of cold 5 mM Tris-HCl buffer, pH 7.6. Subsequently, after short spans of intermittent vortexing and incubations on ice and a 10 min centrigugation at 2310Xg the resultant remnant membrane pellet was suspended in Tris buffer of the same pH and sonicated multiple times with occasional incubations on ice. Following a final 30 min centrifugation at 4390Xg, the supernatant which contained the leishmanial antigen (LAg) was carefully harvested and stored as frozen aliquots until further use in ELISA. For ELISA, plates were first coated with 20μg/ml of LAg in PBS and left overnight at 4°C. Subsequently, after several washes with PBS-0.05% Tween 20, blocking with 1% BSA and further washes at room temperature, mouse plasma samples (1:100 dilution in PBS with 1% BSA) were added to the different antigen coated well and incubation was continued overnight at 4°C. After routine washing, incubation with HRP conjugated anti-mouse IgG goat polyclonal antibody (1: 10,000 dilutions in PBS) was continued for about 3 hrs in room temperature. TMB substrate was applied after washing for color development and following the addition of stop solution, reading was taken at 450 nm using ELISA reader.

### Estimation of Wnt5A in mouse plasma

The level of Wnt5A was estimated from mouse plasma samples. 100µl of 1:10 diluted plasma samples were plated on 96 well polystyrene ELISA plates overnight at 4°C. After 3X wash with PBST, plates were blocked with 1% BSA (Bovine Serum Albumin) for 2 hours. Then the plates were probed with anti-human/mouse Wnt5A antibody (1:1000 dilution) raised in rat. After overnight incubation at 4C, plates were washed with PBST following which incubation with HRP conjugated anti-Rat IgG (1:4000 dilution) for 2 hours was performed. Subsequently TMB (HRP substrate) was added and Stop solution (2N H_2_SO_4_) was used after 30 minutes to stop the colour reaction. Reading was taken in 450 nm in an ELISA plate reader.

### Estimation of Wnt5A in human plasma

Human samples were collected from Murarai village, Birbhum. Samples were tested for rK39 in Malaria Clinic, Calcutta School of Tropical Medicine. For estimating the Wnt5A level in human plasma samples, 96 well ELISA plates were coated with plasma samples (1:50 dilution) overnight at 4°C. After 3X wash with PBST, plates were blocked with 1% BSA for 2 hours. Then the plates were incubated overnight at 4°C with anti-human/mouse Wnt5A antibody (1:2000 dilution) raised in rat. After washing with PBST incubation with HRP conjugated anti-Rat IgG (1:4000 dilution) was continued for 2 hours. Subsequently TMB was added for developing color and Stop solution (2N H_2_SO_4_) was used after 30 minutes to stop the colour reaction. Reading was taken in 450 nm in an ELISA plate reader.

### Estimation of ROS

The fluorescent probe H2DCF-DA was used to check accumulated ROS production in the spleen of Wnt5A+/+ and Wnt5A+/-mice. Single cell suspension was made from the spleens harvested from Wnt5A+/+ and Wnt5A+/-mice. Cells were incubated with H2DCF-DA (10mM) for 20 minutes at 37°C in a CO2 tissue culture incubator. Subsequently, the cells were washed three times with PBS and data acquisition was done in BD.LSR Fortessa Cell analyser. Data was analyzed using FCS express 5 software.

### Statistical analysis

The data were analyzed by Graph Pad Prism 5 software using unpaired t test. Scatter plots and bar graphs are expressed as mean ± SEM. *p* ≤ 0.05 was considered statistically significant. Significance was annotated as follows: \**p* ≤ 0.05, \*\**p* ≤ 0.005, \*\*\**p* ≤ 0.0005.

### Ethics Statement

The use of animals was approved by the Animal Ethics Committee of IICB. The identification number of the approved projects are IICB/AEC/Meeting/2016/Aug, IICB/AEC/Meeting/Aug/2018/4 and IICB/AEC/Meeting/Sep/2019/1. Use of blood from human volunteers was approved by the Ethical Committee of Human Subjects (dated February 10, 2018).

## Results

### Disease from L. donovani infection is more pronounced in Wnt5A heterozygous mice than in the wild type counterparts

In view of the fact that Wnt5A signaling antagonizes *L. donovani* infection in macrophages (19) we wanted to examine if Wnt5A signaling regulates the progression of experimental visceral leishmaniasis. Accordingly, we evaluated the effect of genetic depletion of Wnt5A on the progression of the disease using a mouse model. In essence, we assessed the intensity and outcome of *L. donovani* infection in Wnt5A heterozygous (Wnt5A +/-) mice of B6:129S7 origin (https://www.jax.org/strain/004758), as compared to the wild type (Wnt5A +/+) counterparts. Validation of genetic depletion of Wnt5A in the Wnt5A heterozygous mice and the decreased level of the Wnt5A protein therein as compared to that in the wild type is depicted in Figure S1.

Estimation of *L. donovani* infection in liver and spleen: Leishman Donovan Units (LDU) enumerated from imprints of infected liver and spleen of both sets of mice sacrificed either 45 days or 110 days after infection with either AG83 (antimony sensitive *L. donovani* strain) or BHU575 (antimony resistant *L. donovani* strain) revealed that the liver and spleen of the Wnt5A heterozygous mice harbor significantly greater number of parasites as compared to those of the wild type controls (Figure 1, Panel A). Augmented parasite abundance in the heterozygous mice as compared to the wild type was corroborated by light microscopy of the Giemsa-stained imprints of the infected liver and spleen (Figure 1, Panel B). Consistent with published literature (26), infection in the liver tended to be resolved with the passage of time through 110 days. However, infection prevailed in the spleen, especially in most of the Wnt5A heterozygous mice even after 110 days. As demonstrated by TEM, parasitophorous vacuoles were more frequently visible (shown by red arrowhead) in the spleens of Wnt5A heterozygous mice as compared to those of the wild type (Figure 1, Panel C).

**Figure 1:**
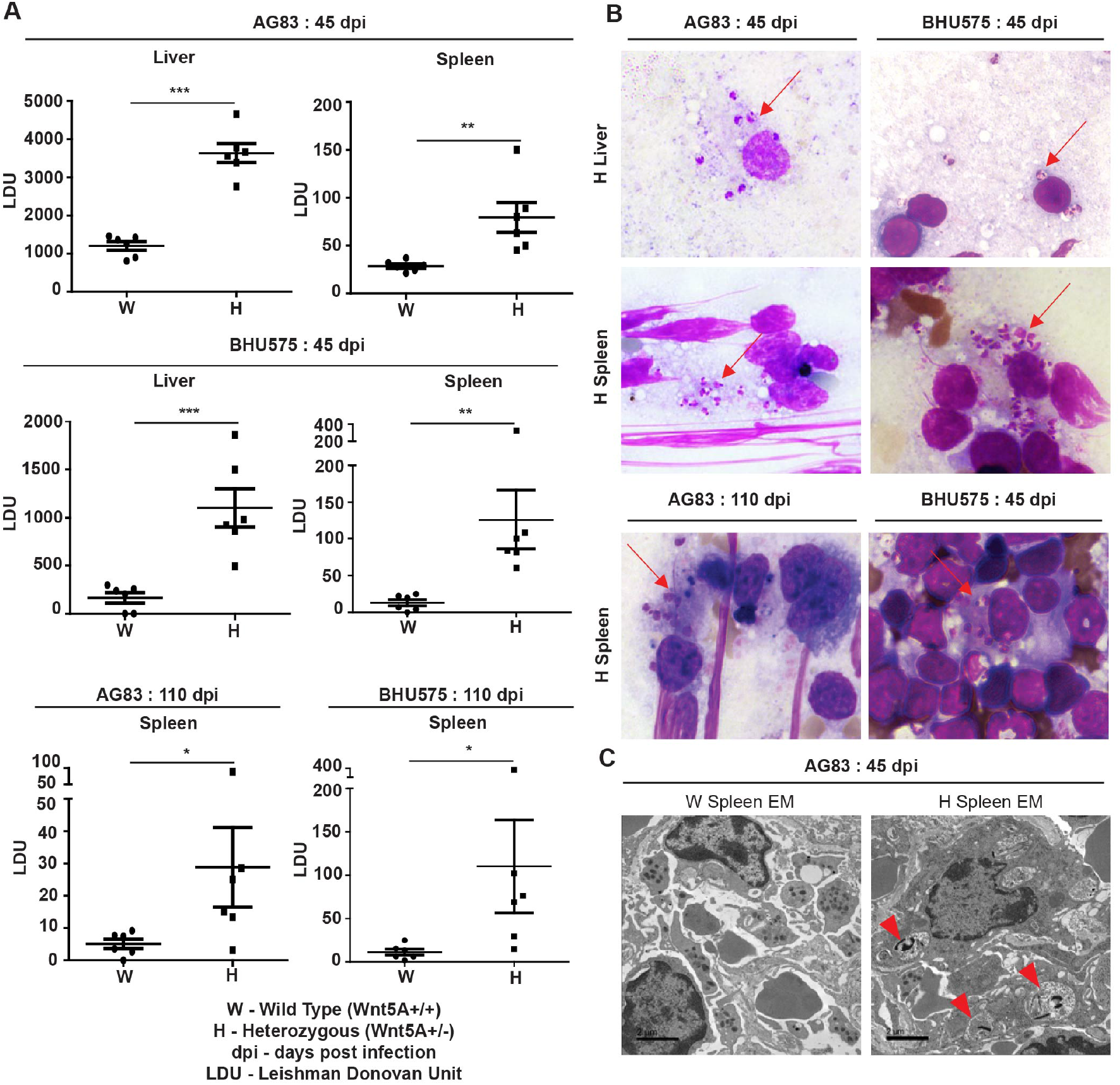
Evaluation of *L. donovani* infection in the liver and spleen of Wnt5A+/+ wild type (W) and Wnt5A+/-heterozygous (H) mice. Microscopic examination of imprints of liver and spleen from infected mice reveals significantly higher parasite load as measured by Leishman Donovan Unit (LDU) in the H category of mice than the control W category, both 45 days and 110 days after infection (dpi) with either the AG83 or BHU575 strain of *L. donovani* (Panel A). Panel B: Representative Giemsa stained micrographs of imprints (under 100X objective/oil immersion) demonstrating *L. donovani* infection with arrowheads. Panel C: TEM micrographs showing parasitophorous vacuoles (PV) in a spleen section of H but not W category after infection. PV was more frequent in H than in W. Data are presented as mean ± SEM. Significance was annotated as follows: \**p* ≤ 0.05, \*\**p* ≤ 0.005, \*\*\**p* ≤ 0.0005. Nuc: hepatocyte nucleus; Mac: macrophages.

Spleen pathology: Increased level of *L. donovani* infection in the Wnt5A heterozygous mice as compared to the wild type correlated with greater severity of disease as revealed by the prevalence of significantly more disrupted splenic germinal centers in the heterozygous mice in comparison to the wild type mice through tissue histology. Upon careful scoring of the levels of organization of splenic germinal centers (clearly explained Materials & Methods), we noted that while the number of organized germinal centers was always significantly more in the wild type mice than in the heterozygous mice, the number of moderately to intensely disorganized germinal centers was always more in the heterozygous mice than in the wild type (Figure 2, Panels A & B). The difference in the numbers of slightly disorganized germinal centers between the two sets of mice was not found to be significant. Higher numbers of CD138+ plasma cells in the spleens of the heterozygous mice, which correlated with increased level of circulating *L. donovani* antigen reactive gammaglobulin (IgG) furthermore, confirmed the disease intensity, as consistent with contemporary literature (Figure 2, Panels C&D) (25, 27). The average weight of spleen per unit body weight was also considerably more in the Wnt5A heterozygous mice following infection with either AG83 or BHU575, as compared to the control, confirming increased infection induced splenomegaly therein (Figure S2).

**Figure 2:**
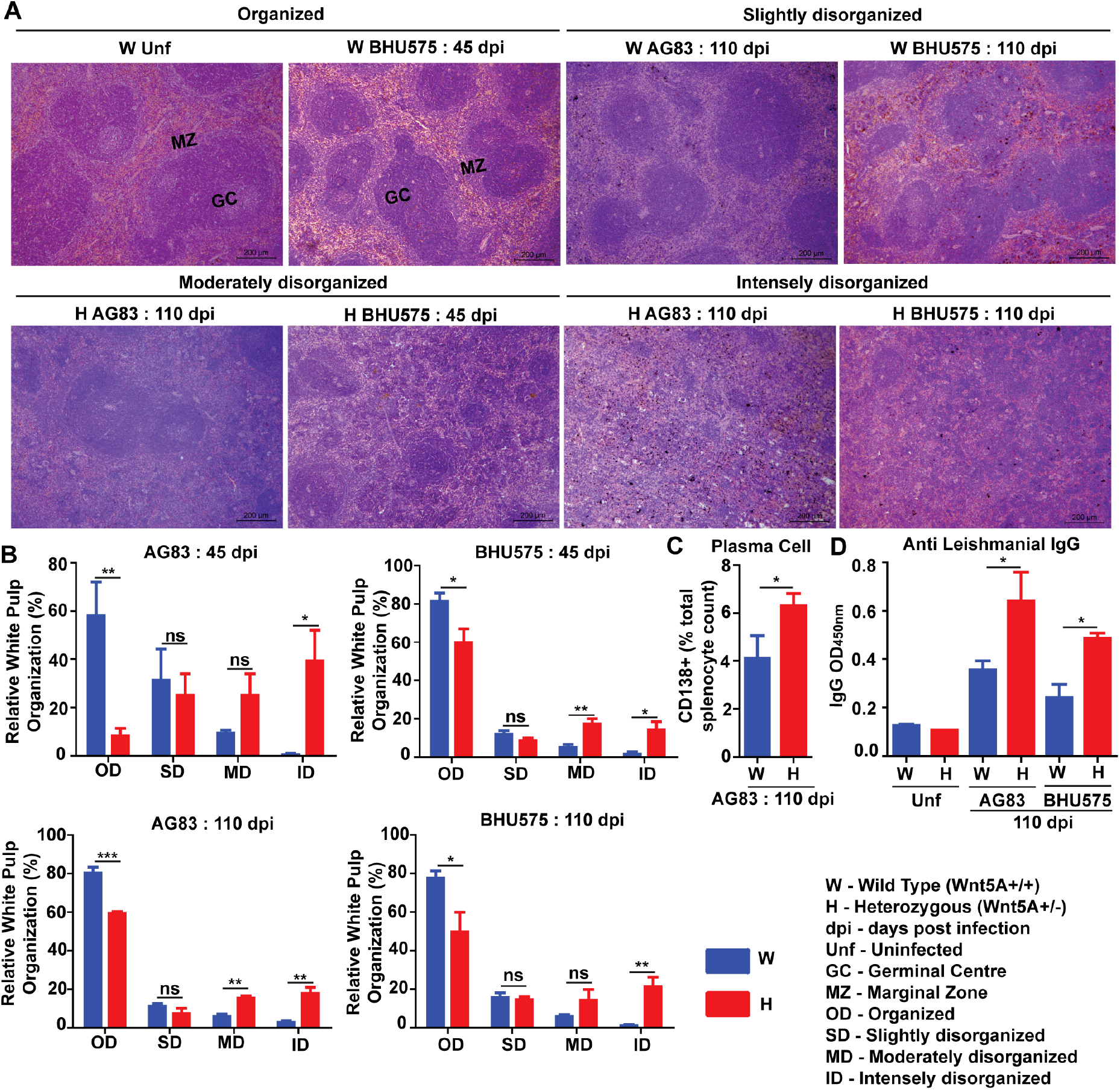
Assessment of spleen pathology in Wnt5A+/- (H) in comparison to Wnt5A+/+ (W) mice after *L. donovani* infection. Panel A: Micrographs (under 10X objective) of representative Hematoxylin & Eosin stained spleen sections from W and H categories, both before and after *L. donovani* infection (45 and 110 dpi), depicting different levels of disorganized splenic white pulp as compared to organized (explained in Materials and Methods). Panel B: Graphical representation of different levels of splenic white pulp disorganization in H mice as compared to W, following infection with either AG83 or BHU575, 45 and 110 dpi. Panels C and D: Graphical representation of higher percentage of plasma cells (C) and increased levels of IgG (D) in H as compared to W, 110 dpi. Data are presented as mean ± SEM. Significance was annotated as follows: \**p* ≤ 0.05, \*\**p* ≤ 0.005, ns: not significant.

Liver pathology: Transmission electron micrographs of liver sections generated 45 days after *L. donovani* infection revealed increased infiltration of monocyte derived macrophages with consequent liver hyperplasia in the Wnt5A heterozygous mice as compared to the wild type controls (Figure 3, Panels A & B), which is in compliance with the correlation of monocyte/macrophage infiltration with active disease (28,29). Augmented disease severity in the Wnt5A heterozygous mice as compared to the wild type was furthermore indicated by the increased prevalence of liver granuloma 45 days after *L. donovani* infection as revealed by histology (Figure 3, Panels C & D) (26).

**Figure 3:**
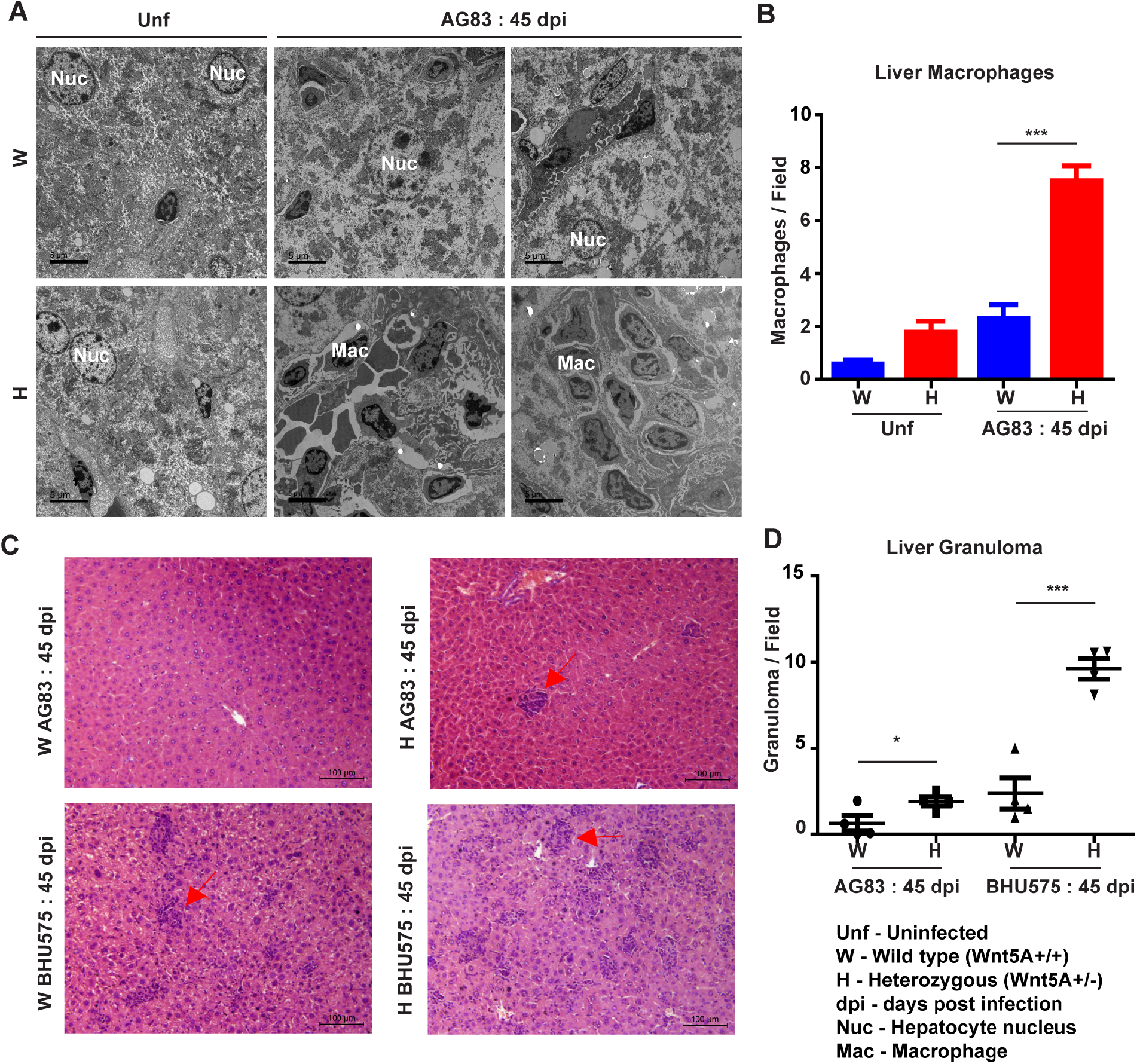
Evaluation of liver pathology of H and W mice after *L. donovani* infection by transmission electron microscopy. Panels A and B: TEM micrographs (A) and morphometric analysis (B) demonstrating increased numbers of monocytes/macrophages in the liver of H as compared W, 45 dpi. ‘Uninfected’ from each category is used as reference. Panels C and D: Similar representation of liver granuloma (under 20X objective) in H as compared to W, 45 dpi. Data from 20 fields are presented as mean ± SEM. Significance was annotated as follows: \**p* ≤ 0.05, \*\*\**p* ≤ 0.0005.

Overall, our results based on careful evaluation of infected liver and spleen imprints, tissue sections and electron micrographs indicated that both the antimony sensitive AG83 and the antimony resistant BHU575 *L. donovani* strains are more infective in Wnt5A depleted condition. We observed that Wnt5A heterozygous mice developed more intense disease after *L. donovani* infection as compared to the wild type controls, where infection was relatively mild. These observations led us to investigate if revamping Wnt5A signaling prevents the development of experimental visceral leishmaniasis.

### Administration of recombinant Wnt5A (rWnt5A) restrains *L*. *donovani* infection and the progression of disease caused by it

In order to evaluate the influence of Wnt5A signaling on the development of experimental visceral leishmaniasis in mice, we examined if the administration of recombinant Wnt5A (rWnt5A) is efficacious in preventing *L. donovani* infection and the progression of disease therein. For our disease model, rWnt5A (100 ng protein in 50μl PBS) was administered intravenously through the tail vein of BALB/c mice on two consecutive days before infection with 10^8^ *L. donovani* promastogotes (either AG83 strain or BHU575 strain) through the same route, and PBS was used as the vehicle control for rWnt5A. Administration of rWnt3A was considered as an additional negative control for this study. All infected mice pretreated separately with rWnt5A, rWnt3A or vehicle control were sacrificed either 45 days or 110 days post infection for enumeration of liver and spleen LDU, and evaluation of liver and spleen architecture by histology, following the same procedures as described before. In compliance with the results obtained with Wnt5A heterozygous and wild type mice, we found that prior injection of rWnt5A but not rWnt3A or just the vehicle control in BALB/c mice inhibits the both the degree of infection with *L. donovani* and the progression of disease. As demonstrated in Figure 4 (Panel A) both liver and spleen LDU were on average considerably less in the rWnt5A treated mice as compared to the PBS and rWnt3A administered controls, both in case of AG83 and BHU575 infection. Giemsa stained representative imprints of infected liver and spleen used for LDU evaluation are depicted in Panel B of the same figure.

**Figure 4:**
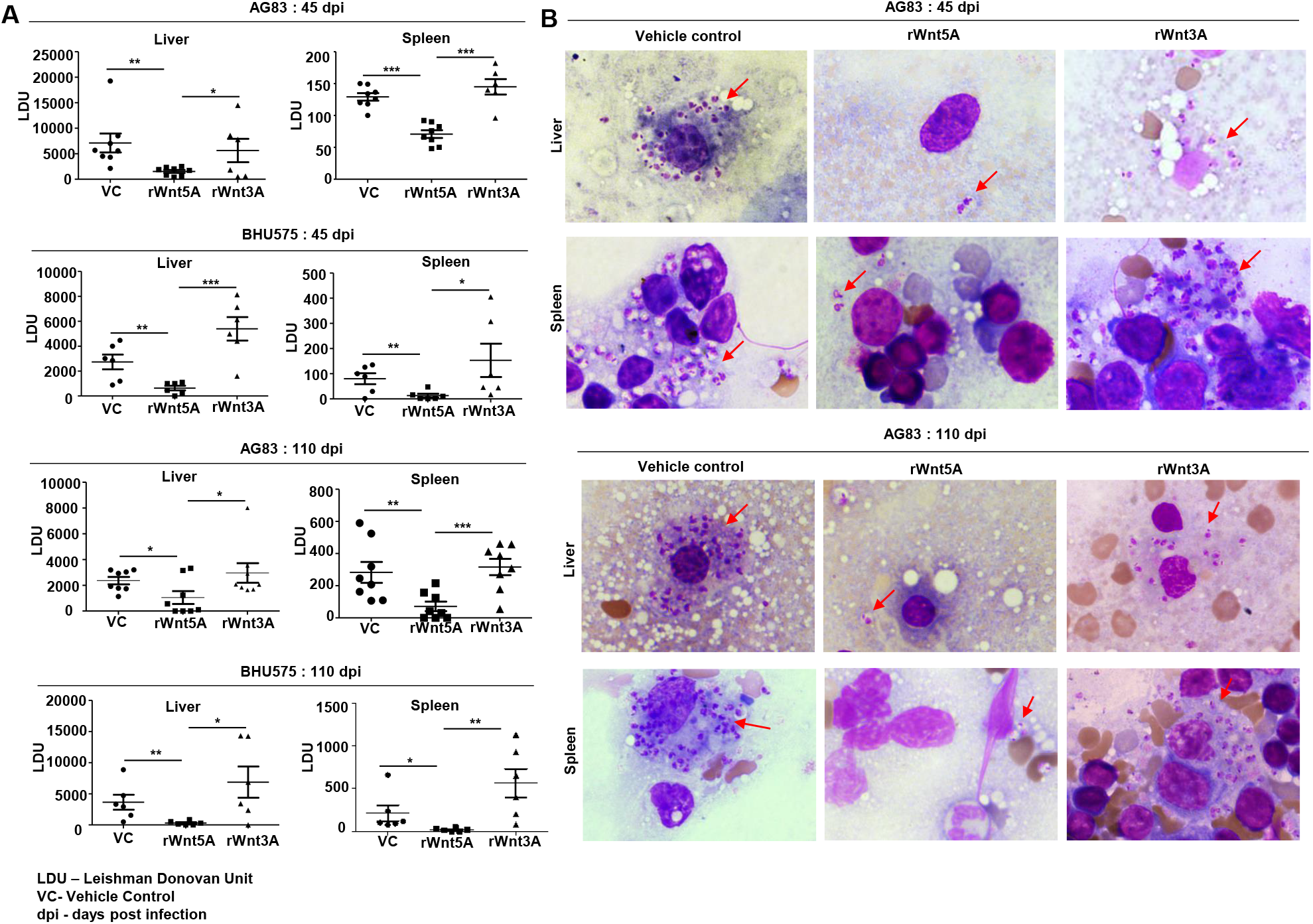
Treatment of mice with rWnt5A restricts *L. donovani* infection. Panel A: Graphical representation of significantly less parasite load (LDU) in spleen and liver imprints of rWnt5A but not rWnt3A pretreated BALB/c mice as compared to corresponding vehicle control (VC: PBS), both 45 and 110 days after infection with either AG83 or BHU575. Panel B: Representative micrographs of Giemsa stained imprints (under 100X objective/oil immersion) of liver and spleen pretreated with rWnt5A, rWnt3A or VC. Data are presented as mean ± SEM. Significance was annotated as follows: \**p* ≤ 0.05, \*\**p* ≤ 0.005, \*\*\**p* ≤ 0.0005.

In agreement with the trend of LDU values obtained from all the splenic imprints, spleens of the *L. donovani* (AG83 or BHU575) infected mice pretreated with rWnt5A revealed significant preservation of germinal center organization much unlike those of mice treated with only PBS or rWnt3A, where marked disorganization of germinal centers was observed. The histological analysis and scoring of germinal center organization in the different sets of mice, which was performed in the same way as described in the previous section is depicted in Figure 5: Panels A & B. Overall, the rWnt5A regimen correlated with the highest numbers of organized and lowest numbers of moderately to intensely disorganized germinal centers as compared to the other two regimens. Significantly lower numbers of CD138+ plasma cells in the spleens of rWnt5A pretreated infected mice as compared to the corresponding controls (PBS and rWnt3A pretreated) matched with the observed intactness of the splenic germinal centers (Panel C). This was in compliance with the relatively low levels of circulating antileishmanial IgG in the rWnt5A treated set (Panel D). Preservation of splenic architecture by rWnt5A treatment was furthermore revealed by blockade of splenomegaly that was induced by *L. donovani* infection (Figure S3). The inhibitory effect of rWnt5A on *L. donovani* infection was also evident from the very low numbers of liver granulomas in the rWnt5A treated infected mice as compared to the controls (PBS and rWnt3A treated) (Figure S4).

**Figure 5:**
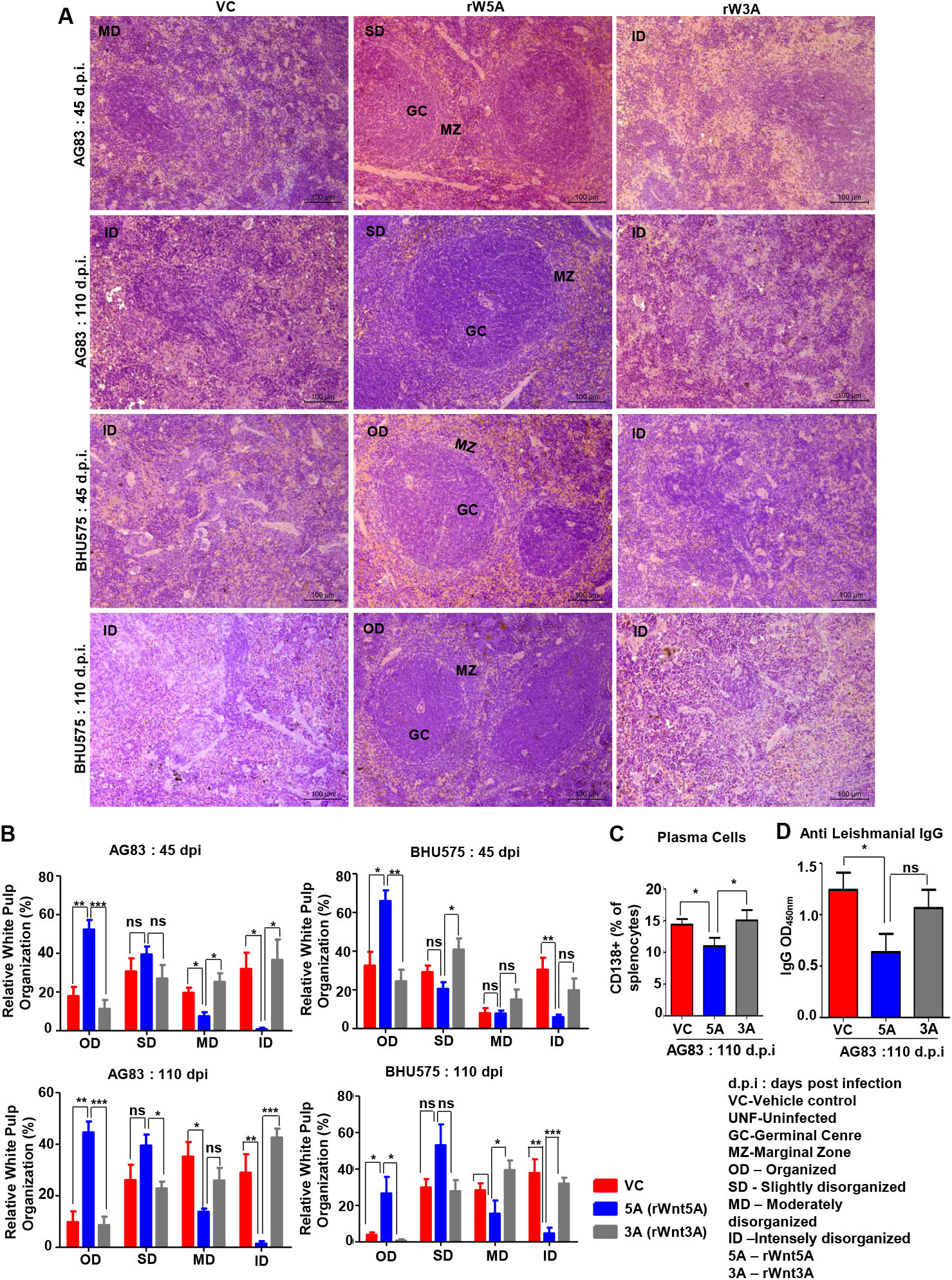
Treatment of mice with rWnt5A inhibits *L. donovani* infection induced disorganization of splenic white pulp. Panel A: Representative micrographs of Hematoxylin & Eosin stained imprints of spleen (under 20X objective) depicting significantly less disorganization in splenic white pulp subject to rWnt5A, but not rWnt3A or VC treatment, prior to AG83 or BHU575 infection. Panel B: Graphical representation of extent of preservation of splenic white pulp organization by rWnt5A treatment before infection as compared to rWnt3A or vehicle control. Panels C & D: Graphical representation of significantly lower percentage of CD138+ plasma cells (C) and anti-leishmanial IgG (D) in rWnt5A pretreated infected mice as compared to those pretreated with rWnt3A or VC. Data are presented as mean ± SEM. Significance was annotated as follows: \**p* ≤ 0.05, \*\**p* ≤ 0.005, \*\*\**p* ≤ 0.0005. ns: not significant.

### Inhibition of progression of experimental visceral leishmaniasis by Wnt5A is associated with an altered profile of macrophages, T cells and secreted cytokines

In view of the documented role of Wnt5A signaling in macrophage and T cell activation (6– 10,19,29,30) we wanted to examine if the Wnt5A mediated inhibition of disease progression in mice by *L. donovani* infection is linked with changes in the status of macrophages, T cells and cytokines. In essence, we evaluated the immune potential of rWnt5A in relation to rWnt3A and the vehicle control (vc), toward restriction of disease progression.

Flow cytometry performed on splenocytes harvested from the different experimental sets of rWnt5A/rWnt3A/vc pretreated *L. donovani* infected BALB/c mice revealed that the rWnt5A pretreated mice harbored on average higher numbers of activated splenic CD3+CD4+ and CD3+CD8+ T cells, positive for both intracellular IFN-γ and granzyme B (GrB) as compared to those pretreated with just vc or rWnt3A (Figure 6, Panel A). PBMC harvested from these mice exhibited a similar trend in the level of CD3+CD4+ and CD3+CD8+ T cells (Figure 6, Panel B). In concurrence with these findings, we observed that the percentage of CD3+CD4+ and CD3+CD8+ T cells in *L. donovani* infected Wnt5A heterozygous mice was considerably less than that in the corresponding wild type controls (Figure 6, Panel C).

**Figure 6:**
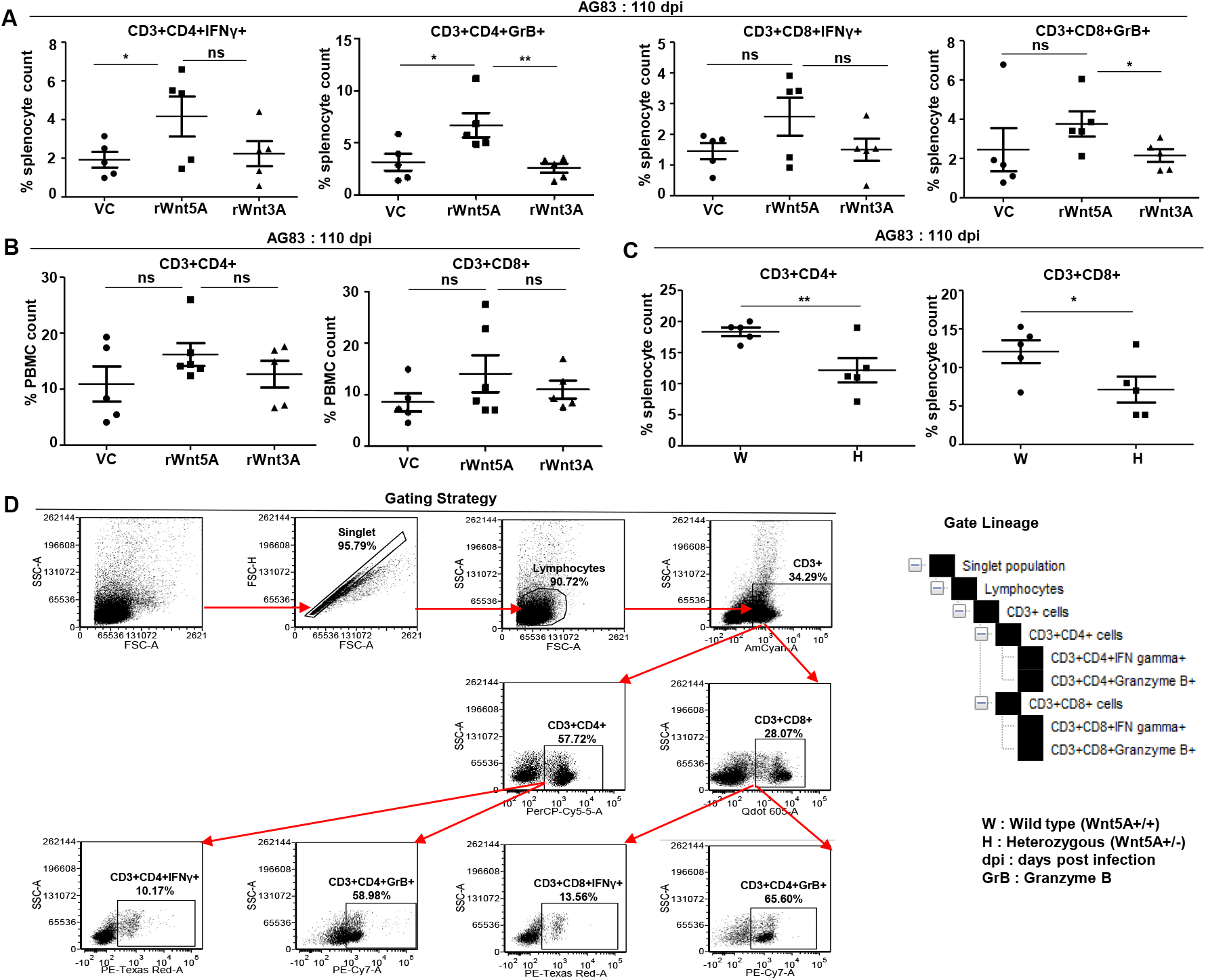
Wnt5A supports CD4 and CD8 T cell maintenance and activation in the backdrop of *L. donovani* infection. Panel A: Graphical representation of increased percentage of IFN*γ* positive and GrB positive splenic CD3+CD4+ and CD3+CD8+ T cells subject to rWnt5A treatment before infection, in comparison to the corresponding controls, as depicted by FACS. Panel B: Representation of a similar trend in the PBMC harvested from the same sets of mice. Panel C: Graphical representation of lesser percentages of splenic CD3+CD4+ and CD3+CD8+ T cells in *L. donovani* infected H mice as compared to the controls. Panel D: Depiction of FACS gating strategy. Data are presented as mean ± SEM. Significance was annotated as follows: \**p* ≤ 0.05, \*\**p* ≤ 0.005, ns: not significant.

In association with our observation of the Wnt5A dependent prevalence of splenic CD4 and CD8 T cells in the backdrop of *L. donovani* infection, we demonstrated that Wnt5A pretreatment of *L. donovani* infected mice led to a moderately increased presence of CD169+ marginal zone metallophillic macrophages (MMM), which are known to engulf pathogens, and collaborate with white pulp dendritic cells for antigen presentation to T cells (Figure 7, Panel A) (31,32). Additionally, as shown by confocal microscopy, Wnt5A associated MMM abundance was coupled with a structured occurrence of CD209+ marginal zone (MZ) macrophages, also known for their ability to internalize and destroy pathogens, (Figure 7, Panel C) (33). Such profile of MMM and MZ macrophages was not observed in case of pretreatment of the infected mice with rWnt5A or vc (Panels A & C). In both vc and rWnt3A treated sets, there were relatively lesser numbers of CD169+MMM and absence of a structured organization of CD209+MZ. Furthermore, as demonstrated by flow cytometry rWnt5A pretreated mice infected with *L. donovani* also harbored increased numbers F4/80+ red pulp macrophages, known for their microbe scavenging property (33), in comparison to the vc and rWNT3A pretreated controls (Figure 7, Panel B).

**Figure 7:**
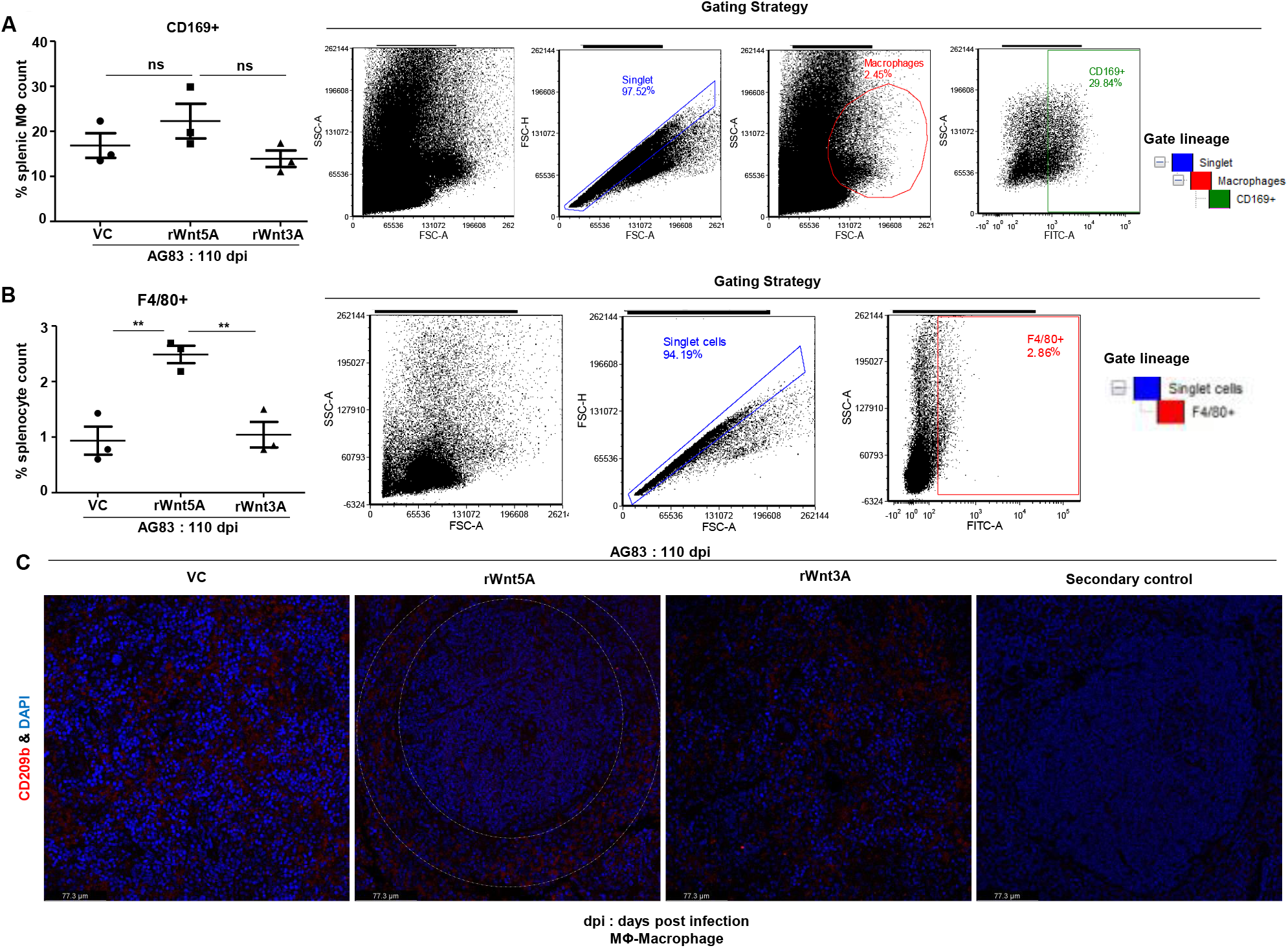
rWnt5A supports preservation of splenic macrophages after *L. donovani* infection. Panel A: Graphical representation of increased percentage of CD169+ splenic marginal metallophillic macrophages associated with rWnt5A treatment as compared to controls, with outline of FACS gating strategy. Panel B: Increased percentage of F4/80+ macrophages in the same sets, with outline of FACS gating. Panel C: Preservation of CD209+ splenic marginal zone macrophages (denoted by white dashed concentric lines) in association with treatment of rWnt5A, but not rWnt3A or VC, as revealed by confocal microscopy (under 40X objective/ oil immersion). Data are presented as mean ± SEM. Significance was annotated as follows: \*\**p* ≤ 0.005, ns: not significant.

Wnt5A signaling supported the making of a TH1 like cytokine profile, which is known to antagonize *L. donovani* infection (34) in conjunction with the preservation of splenic T cells and macrophages. Evaluation of the cytokine profile of blood plasma of both Wnt5A+/+ and Wnt5A+/-mice infected with *L. donovani* (either AG83 or BHU575) revealed considerably higher IFN*γ*/IL10 ratio in the plasma samples of Wnt5A+/+ mice in comparison to those of the Wnt5A+/-mice (Figure 8, Panel A). Revamping of Wnt5A signaling by rWnt5A treatment before *L. donovani* infection, furthermore, resulted in increase in the plasma IFN*γ*/IL10 ratio. This was not observed when rWnt3A was used as a control instead of rWnt5A (Panel B).

**Figure 8:**
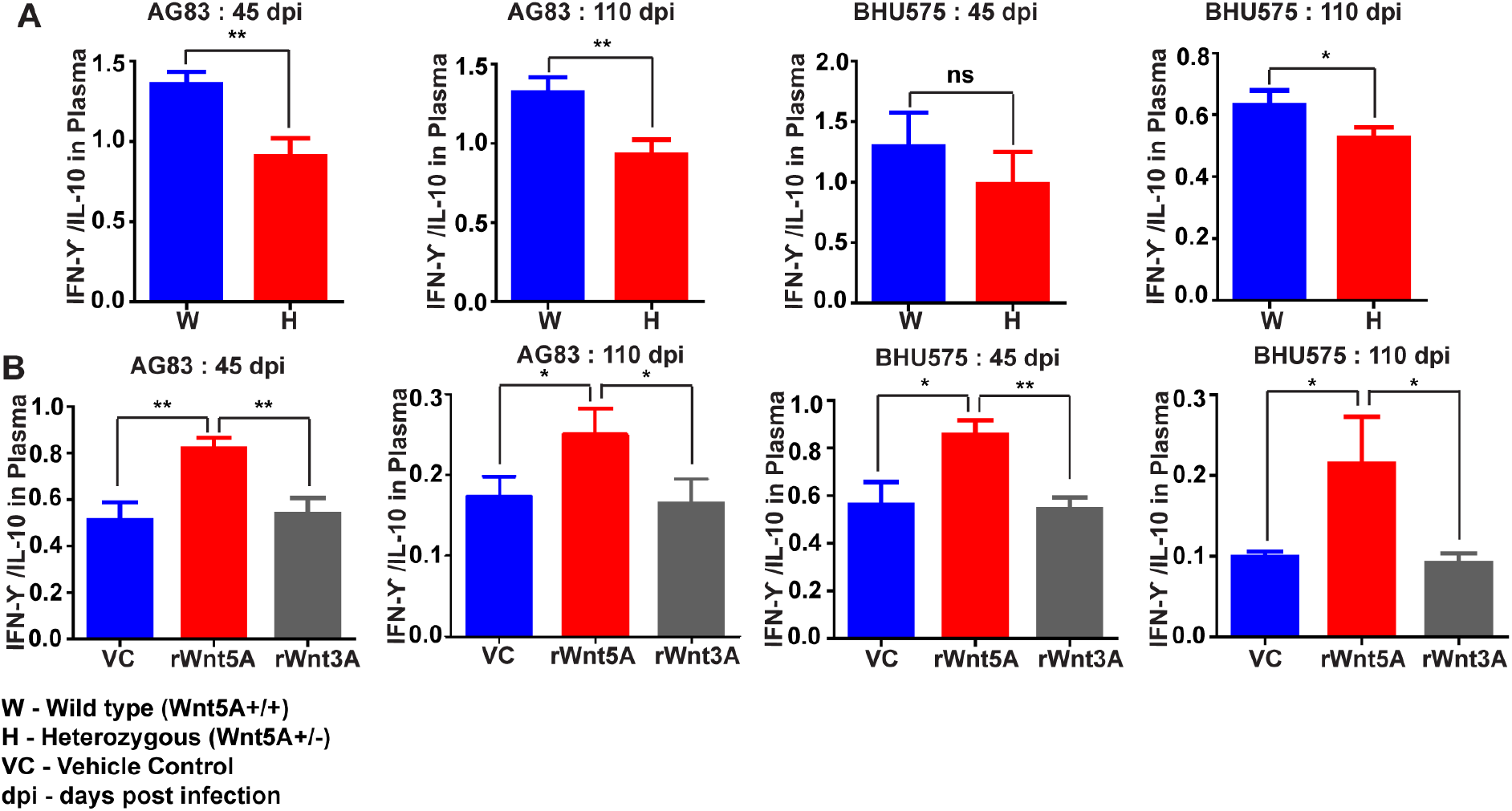
rWnt5A promotes a TH1 like cytokine thrust during *L. donovani* infection. Panel A: Graphical representation of higher IFN*γ*/IL10 ratio in the blood plasma of AG83 and BHU575 infected Wnt5A+/+ (W) mice as compared to Wnt5A+/- (H) mice. Panel B: rWnt5A treatment before AG83 or BHU575 infection correlates with higher plasma IFN*γ*/IL10 in comparison to rWnt3A or VC treatment. Data are presented as mean ± SEM. Significance was annotated as follows: \**p* ≤ 0.05, \*\**p* ≤ 0.005, ns: not significant.

Altogether, these results suggest that the altered status of splenic macrophages and T cells, and the TH1 like cytokine thrust associated with Wnt5A signaling support parasite clearance and help restrict disease progression in experimental visceral leishmaniasis. The augmented level of Wnt5A dependent ROS in the spleens of infected mice (Figure S5) is in line with the altered splenic milieu and parasite killing linked with Wnt5A signaling (1,26,35).

## Discussion

In view of the fact that Wnt5A signaling antagonizes *L. donovani* infection in macrophages (19), we evaluated the potential of Wnt5A signaling in restraining the progression of disease in experimental visceral leishmaniasis. In this study we demonstrated that while depletion of Wnt5A in Wnt5A heterozygous mice increases susceptibility to experimental visceral leishmaniasis, administration of rWnt5A helps to contain the disease by restraining *L. donovani* infection and disease progression, as clearly depicted in Figures 1 – 5. The observed association of the increase in macrophage numbers, CD4/CD8 T cell activation and TH1 like cytokine thrust with Wnt5A mediated blockade in disease progression in experimental animals (Figures 7 & 8) indicates that parasite clearance by macrophages and T cells is facilitated through Wnt5A signaling at the onset of infection.

Previous findings from *in vitro* studies indicated that Wnt5A induced alteration of actin dynamics contributes to the reduction of parasite burden in *L. donovani* infected macrophages (19). Figure S6 explicitly depicts the disruption of supposedly protective actin ring niches that are usually present around the parasite containing vacuoles in *L. donovani* infected peritoneal macrophages, through activation of Wnt5A signaling. Such remodeling of cytoskeletal actin in the macrophages of the red pulp (F4/80+) and white pulp (CD169+ and CD209b+), which are preserved by Wnt5A signaling in infected animals (Figure 7), could lead to inhibition in actin niche formation around parasitophorous vacuoles and destruction of parasites. The marginal zone (CD209b) and marginal metallophillic (CD169) macrophages, moreover could aid in the presentation of dead parasite fragments to T cells leading to T cell activation and cytokine secretion (30–33). Such intercellular coordination during infection may lead to a sustained activation of macrophages and T cells targeted toward parasite clearance.

Overall, our animal experiments indicate that Wnt5A signaling reduces susceptibility to *L. donovani* infection and progression of visceral leishmaniasis. In a very preliminary study comprising only a limited number of blood plasma samples collected from healthy individuals and patients with VL or PKDL (Post Kalazaar Dermal Leishmaniasis), we in fact noted, that healthy individuals and most of the patients either under treatment with miltefosine or with no symptomatic VL had higher level of Wnt5A in their plasma than those with symptomatic VL or PKDL. (Table1 & Figure S7). It will thus be important to examine the relevance of Wnt5A in relation to VL/PKDL using an appropriate number of healthy volunteers and test subjects with a clear knowledge of the associated history of the disease. Furthermore, future experiments directed toward the use of recombinant Wnt5A as a prophylactic or therapeutic agent may prove useful for restricting progression of the recalcitrant disease that is either resistant or poorly managed by the currently administered drugs.

## Acknowledgements

The authors thank Chandan Bhattacharya for tissue histology, Tanmoy Dalui for FACS analysis, Shounak Bhattacharya for confocal microscopy, CSIR-IICB central instrument facility for instrument support, CSIR-IICB animal house facility for animal breeding & maintenance, and Indrajit Sikder for overall technical assistance. The authors also thank Suranjan Bhattacharya and other members of the Lions Club, Birbhum, WB, India for organizing a camp for the collection of human test samples. This work was supported by a grant from the Indian Council of Medical Research (ICMR), Govt. of India (No. 2016-0222/CMB/ADHOC/BMS) and institutional funding. Shreyasi Maity is supported by a fellowship from the University Grants Commission, Govt. of India. Syamal Roy acknowledges ICMR for the Emeritus Fellowship.

