## Supplemental File for "Wnt5A Signaling Blocks Progression of Experimental Visceral Leishmaniasis"

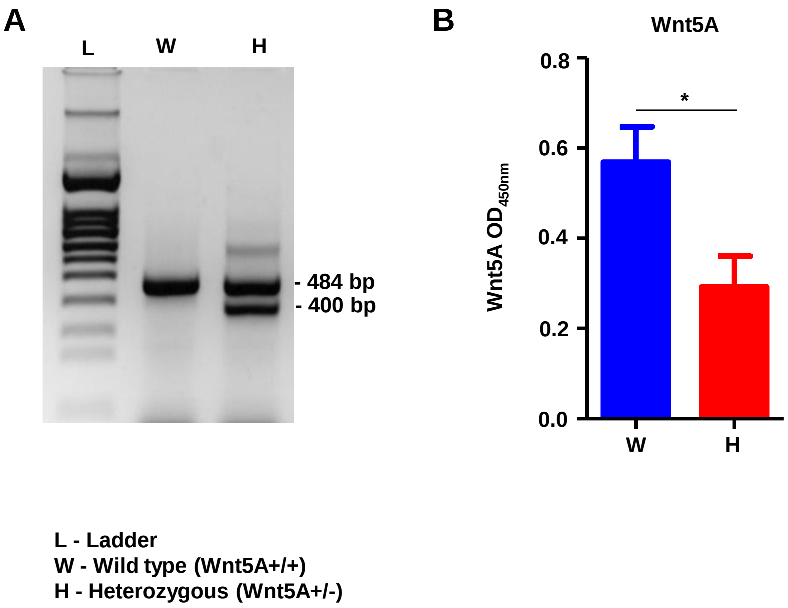


Figure S1: <u>Characterization of Wnt5A heterozygous mice</u>. Panel A: Identification of Wnt5A+/- (heterozygous) mouse by genotyping of DNA extracted from tail by PCR, following Jackson Laboratory protocol. Panel B: Graphical representation of relatively less Wnt5A protein in the plasma of Wnt5A heterozygous mice as compared to wild type, as demonstrated by ELISA.

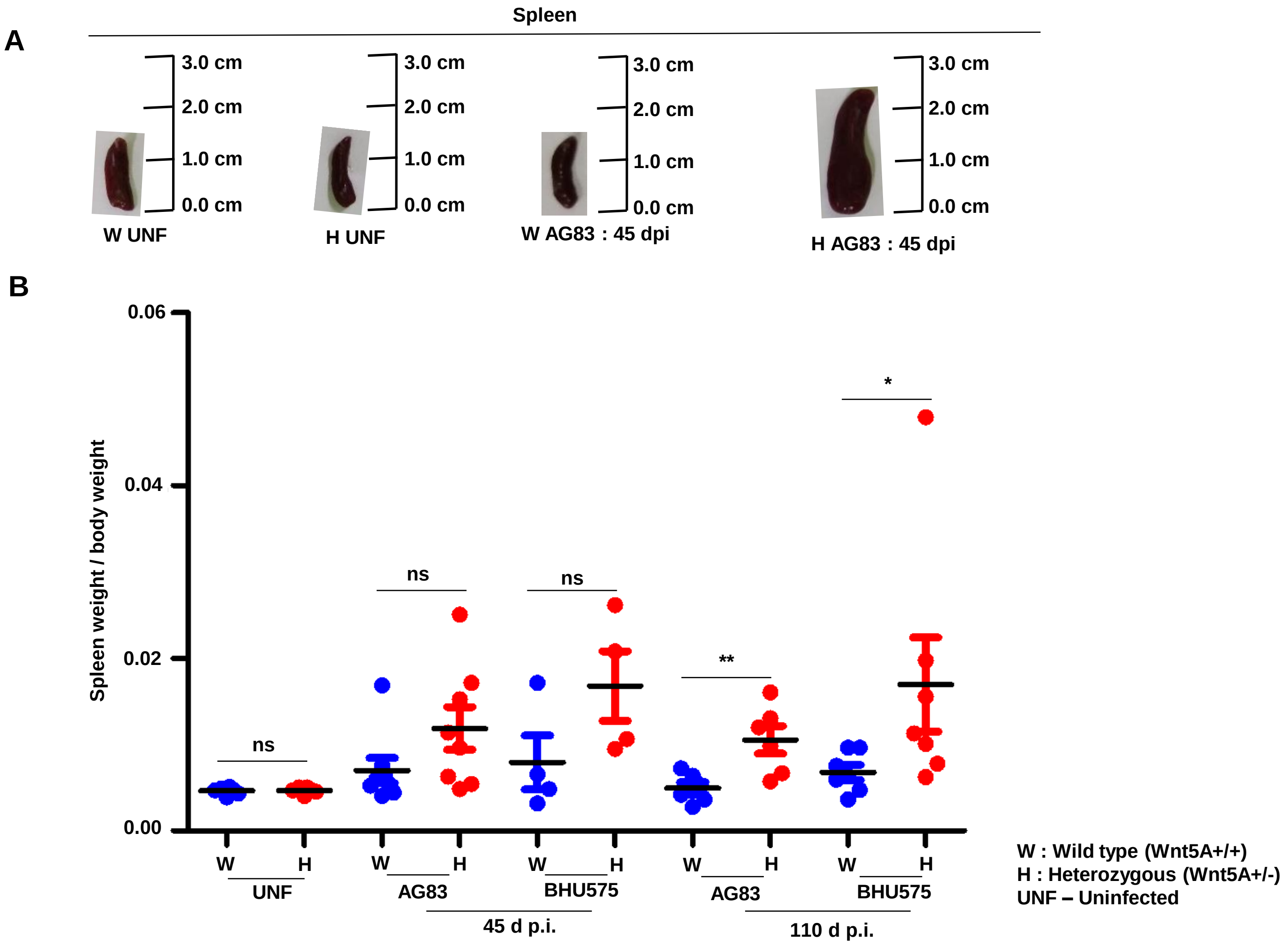


Figure S2: Wnt5A heterozygous (H) mice show increased frequency of splenomegaly as compared to wild type (W) controls upon *L. donovani* infection. Panel A: Representation of calibration of spleen size. Panel B: Graphical representation of spleen weight in the H category as compared to W, after infection. d.p.i: days post infection.

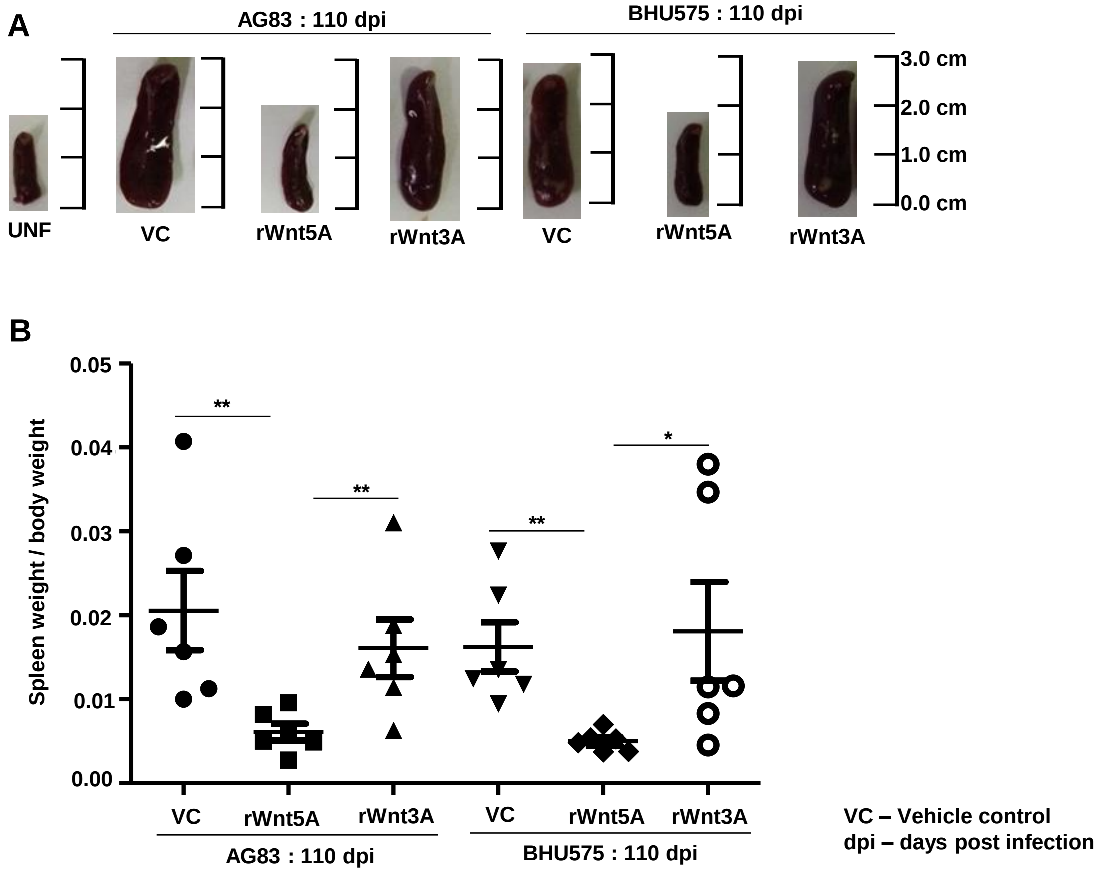


Figure S3: Incidence of *L. donovani* infection induced splenomegaly is lower in rWnt5A treated mice than in the corresponding controls. Panel A: Representation of calibration of spleen size. Panel B: Graphical representation of spleen weight in Wnt5A, Wnt3A or vehicle control (VC) treated mice demonstrating reduced frequency of splenomegaly in rWnt5A treated infected mice as compared to those treated with rWnt3A or VC.

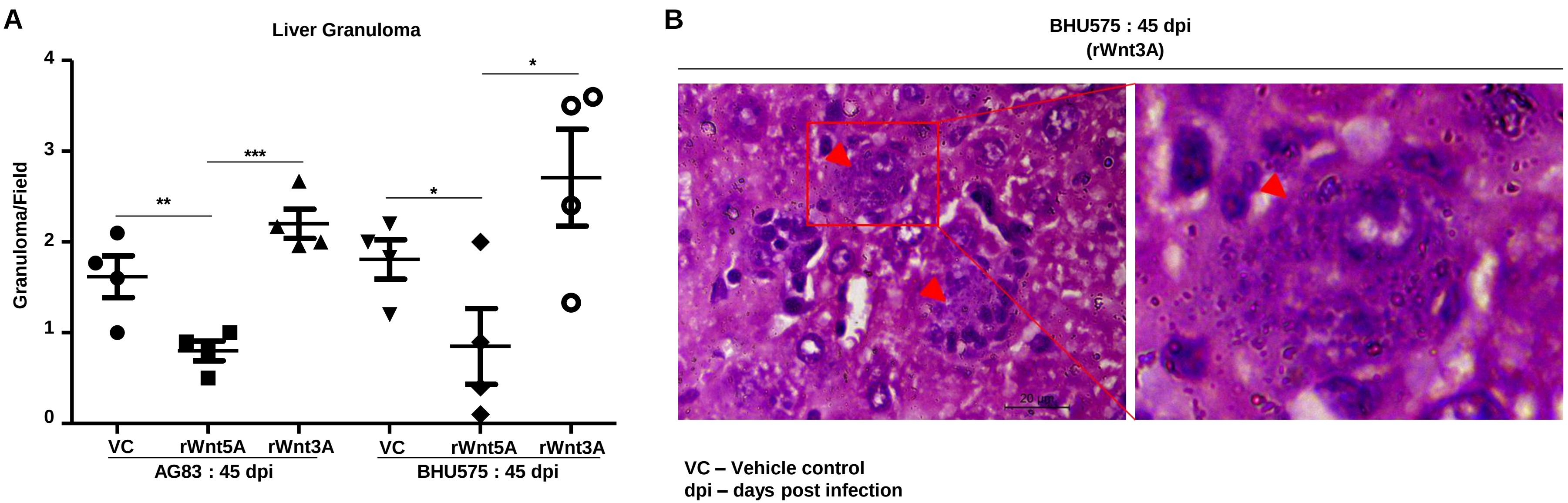


Figure S4: <u>Frequency of liver granuloma is significantly less in *L. donovani* infected mice pretreated with rWnt5A, but not rWnt3A or VC</u>. Panel A: Graphical representation of reduced frequency of liver granuloma subject to rWnt5A pretreatment of infected mice as opposed to pretreatment with rWnt3A or VC. B: Micrograph of H & E-stained liver section (under 100X objective/oil) demonstrating the presence of parasites inside the liver granuloma of an rWnt3A pretreated BHU575 infected mouse.

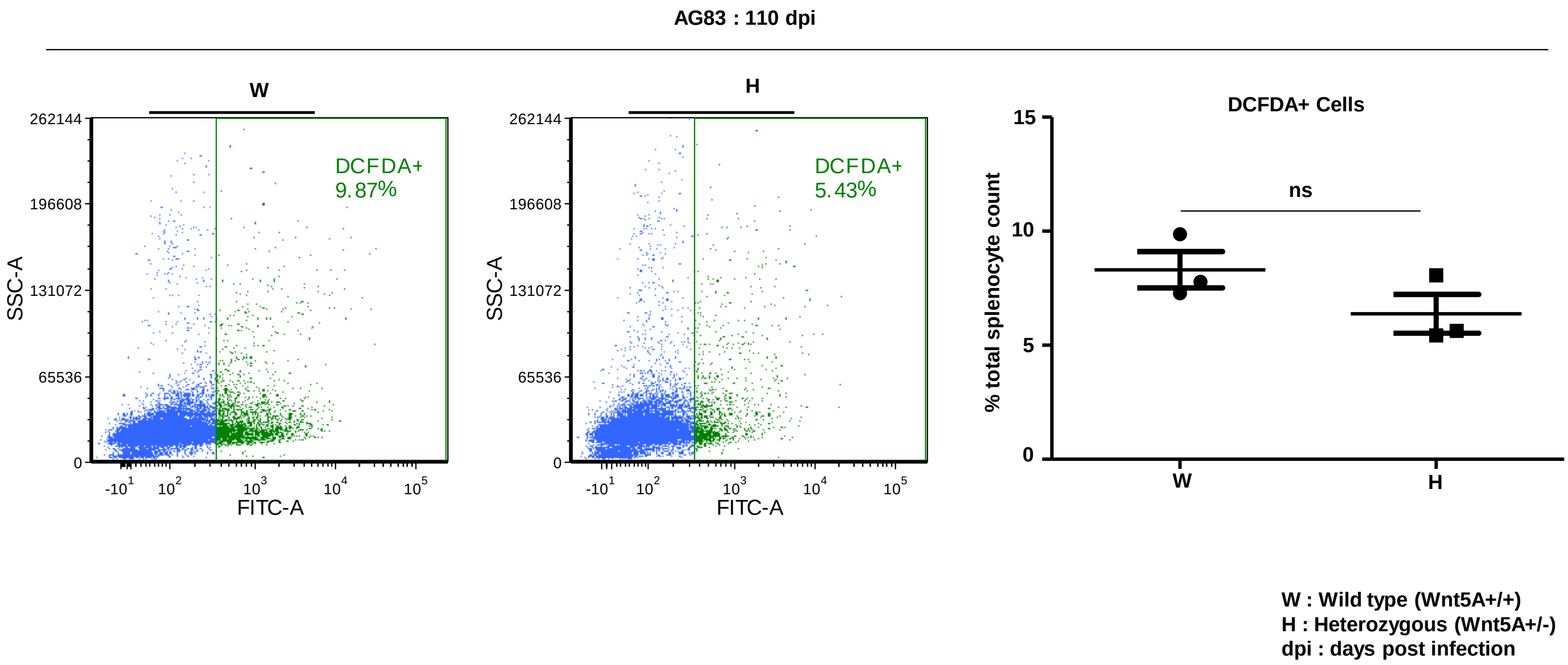


Figure S5: Wnt5A associated restriction of *L. donovani* infection is linked with induction of ROS. Wnt5A+/+ (wild type) *L. donovani* (AG83) infected mice have more ROS positive cells in the spleen in comparison to the Wnt5A+/- (Wnt5A heterozygous) counterparts, as demonstrated by FACS, using DCFDA for ROS detection. Higher ROS correlates with low parasite count (explained in main text).

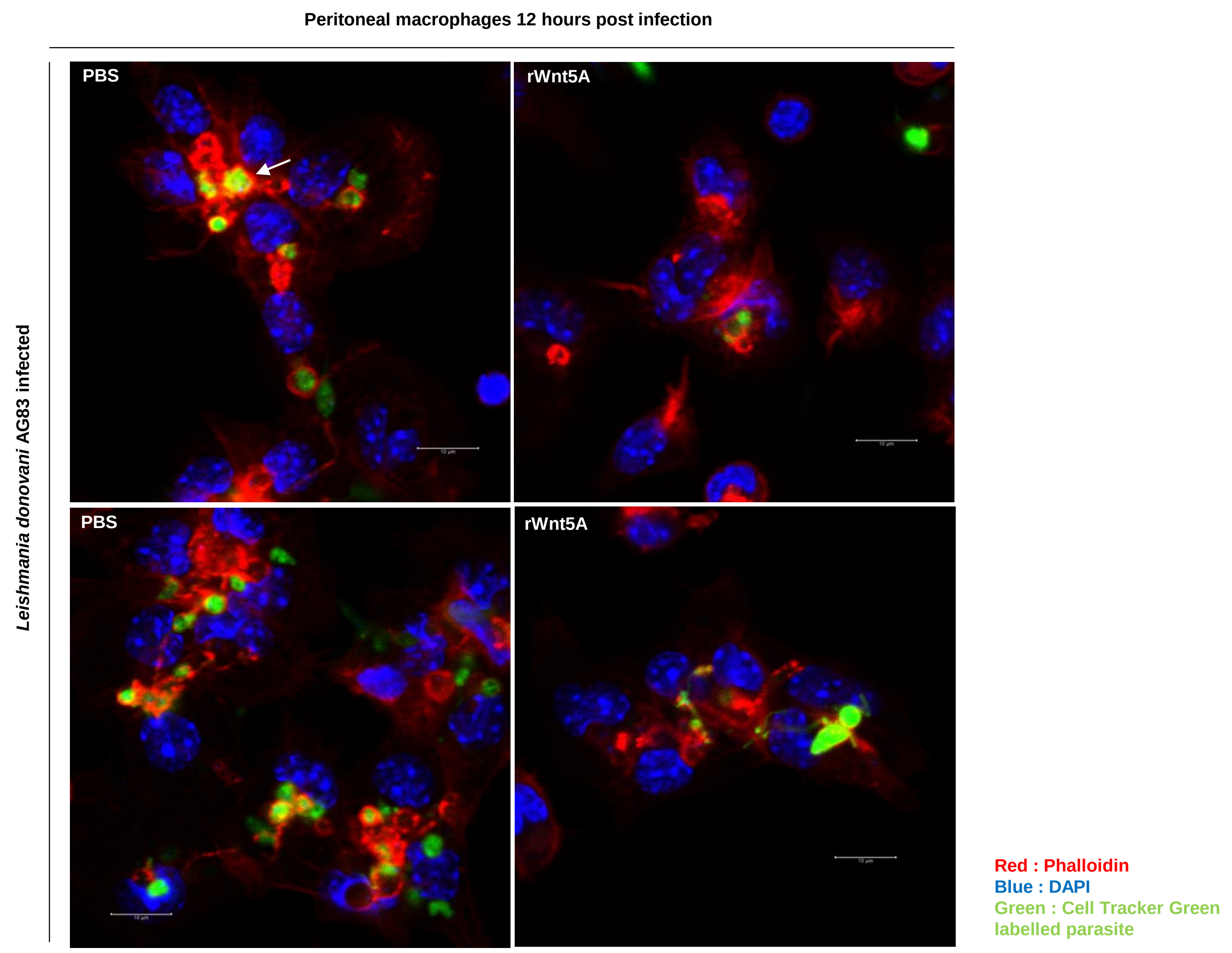


Figure S6: <u>Wnt5A associated restriction of *L.donovani* infection correlates with remodeling of the actin cytoskeleton</u>. Confocal microscopy of phalloidin stained peritoneal macrophages infected with *L. donovani* demonstrates rWnt5A treatment associated disruption of actin rings that are otherwise present after infection. Arrow marks denote actin rings (under 63X objective/oil immersion/2.51 zoom).

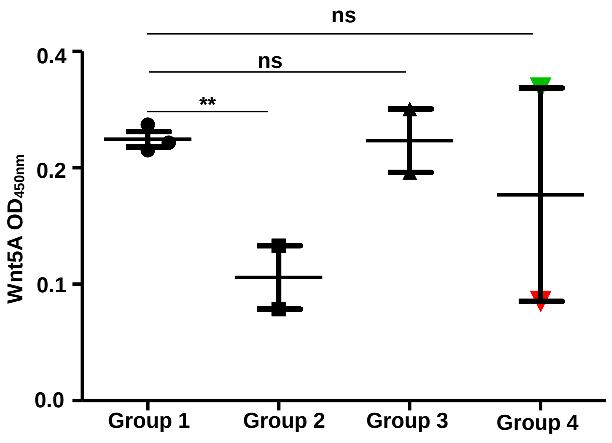


**Group 1 – Healthy Individuals**
**Group 2 – rK39- with lesions**
**Group 3 – rK39+ with lesions and under Miltefosine treatment**
**Group 4 – rK39+ without lesion, no treatment**

Figure S7: Varied level of Wnt5A in the plasma of healthy individuals and those with history of visceral leishmaniasis (VL) or post-kalazar visceral leishmaniasis (PKDL). Graphical representation of varied level of plasma Wnt5A in healthy individuals and those diagnosed with VL/PKDL based on reactivity to rk39 antibody. Patient history is explained in Table 1. Parasite clearance may be associated with the level plasma Wnt5A.

| Sample Number | Past symptoms | Current symptoms | Anti-Leishmanial antibody test (rK39 positve / negative) | Treatment taking |
|---|---|---|---|---|
| Sample 1 **(Group 3)** | | Hypopigmented skin rash all over the body. No fever, no organomegaly | + | Miltefosine for about 2 months 20 days |
| Sample 2 **(Group 2)** | She was diagnosed on 27/07/2018. Skin scraping test was positive for LD bodies. On 27/07/2018, Hb was 7.8 g/dL (at disease onset), WBC count was 7,180. | Treated case with nearly complete recovery. Small lesion was found near her lower lip. | - | Treatment with Miltefosine completed on 2018 |
| Sample 3 **(Group 3)** | VL 7 years ago | Hypopigmented spots over face. No fever, no organomegaly | + | Miltefosine for last 3 months |
| Sample 4 **(Group 4)** | VL 10 years ago | | + | |
| Sample 5 **(Group 1)** | No history of Kalaazar | | - | |
| Sample 6 **(Group 2)** | No history of fever | | - | |
| Sample 7 **(Group 4)** | Treatment completed on July 2018 from Murarai hospital. Before treatment splenomegaly was found as 3 finger, Hb was 5.8g/dL, WBC count was 4,700. | No organomegaly | + | Treatment with Miltefosine completed on July 2018 |
| Sample 8 **(Group 1)** | | Apparently healthy | | |
| Sample 9 **(Group 1)** | | Apparently healthy | | |

Supplementary Table 1: Characteristics of blood samples from human subjects
